# Defining common principles of gene co-expression refines molecular stratification in cancer

**DOI:** 10.1101/372557

**Authors:** Matthew A. Care, David R. Westhead, Reuben M. Tooze

**Affiliations:** Section of Experimental Haematology, Leeds Institute of Cancer and Pathology, University of Leeds, Leeds, LS9 7TF, UK; Bioinformatics Group, School of Molecular and Cellular Biology, University of Leeds

**Keywords:** Cancer Hallmarks, Molecular classification, Breast Cancer, Colon Cancer, Network analysis, Gene expression, Mast cell

## Abstract

Cancers converge onto shared patterns that arise from constraints placed by the biology of the originating cell lineage and microenvironment on recurrent programs driven by oncogenic events. This structure should be transferable to molecular stratification. We exploit expression data resources and a parsimonious and computationally efficient network analysis method to define consistent expression modules in colon and breast cancer. Comparison between cancer types identifies principles of gene co-expression: cancer hallmarks, functional and structural gene batteries, copy number variation and biology of originating lineage. Mapping outcome data at gene and module level onto these networks generates a detailed interactive resource. Testing the utility of the resulting modules in TCGA data defines specific associations of module expression with mutation state, identifying striking associations such as mast cell gene expression and mutation pattern in breast cancer. These analyses provide evidence for a generalizable framework to enhance molecular stratification in cancer.

## Introduction

A primary driver in tumor classification is enhanced precision through molecular characterization. Such analysis provides an increasingly complex view of individual tumor biology [1], resulting in the concept of combinatorial characterization using multiple platforms. An extension is provided by pan-cancer classification where cases associated with key molecular features are combined potentially across the boundaries of conventional classification [2].

Gene expression-based classifications have defined both prognostically and pathogenetically distinct cancer subtypes [3-6], which have preferential association with mutational and cytogenetic profiles [7]. Use of reduced sets of genes allows recognition of subtypes in applied classifications [8, 9]. The cancer hallmark paradigm postulates that aberrantly regulated features assemble in modular fashion to promote malignancy [10]. Thus, an integrated assessment of these features might also take a modular approach within individual cancers.

With multiple data sets the pattern of correlation between individual pairs of genes can be used to determine intrinsic modules of gene co-expression [11]. Exemplifying how modular patterns of co-expression can be identified within the overall profile of a tumor, gene expression allows inference of tumor infiltrating immune populations [12, 13]. Existing expression data sets provide an extensive resource for individual types of cancer, which sit alongside multiparameter analysis across diverse cancer types in resources such as TCGA. Applying a common data-led analysis strategy of gene co-expression to different cancer types, should discover shared modules of expression linked to the neoplastic state between cancer types alongside features of established expression-based classifications. Previous successful methods for network analysis have provided significant insights in model systems and clinical data [14-17]. A challenge in network-based analysis is high density of connectivity, but this can be successfully negotiated using approaches that focus onto modular patterns of gene expression [18]. Here we test a conceptually simple, parsimonious approach to the problem of connectivity reduction, as means to derive modular expression networks and a platform that facilitates linkage between pre-existing gene expression assets and exploration of multi-parameter data such as TCGA.

## Results

### Parsimony enhances gene expression network clustering

We reasoned that a parsimonious approach in which only a restricted number of the most significant correlations (edges) per gene (node) are retained might provide a focusing effect in network analysis. To address this, we developed a method in which only the most highly correlated genes are retained for each index gene. These are assembled into a correlation matrix in which an index gene may re-acquire additional correlations if it represents a common retained partner of other genes in the matrix (Fig. 1A, S1A; full pairwise gene correlation lists online). The resulting parsimonious correlation matrices were used in network generation.

**Figure 1.**
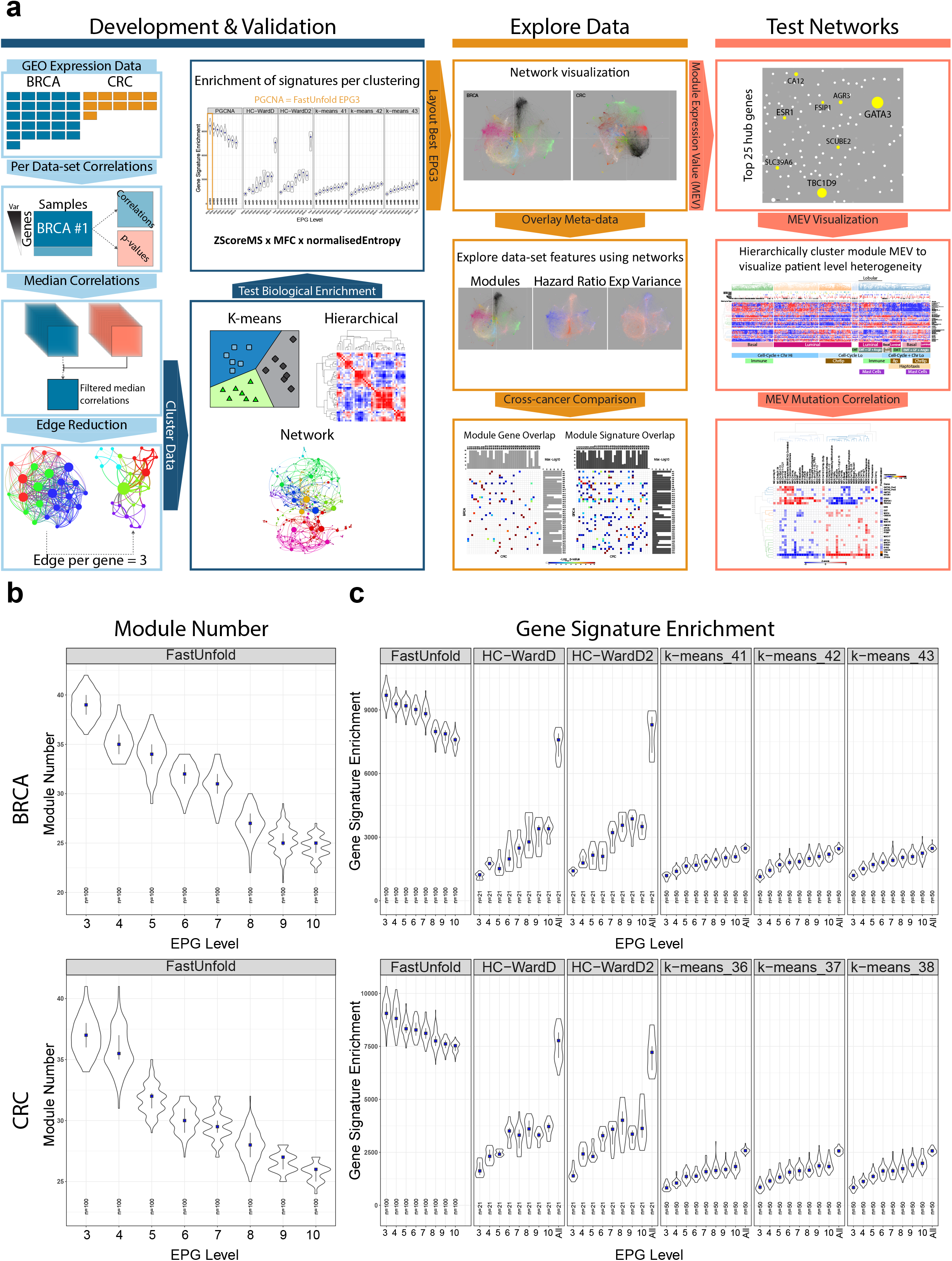
Process outline and validation of parsimony in network generation. **(A)** Graphical outline of the methodological steps. Networks and linked resources available at http://pgcna.gets-it.net/ **(B)** Violin plots of the distribution module numbers from FastUnfold clustering of matrices with different levels of edge reduction (edges per gene EPG). Upper panel BRCA, lower panel CRC. **(C)** Distribution of gene signature enrichment observed with FastUnfold (n=100 runs per EPG), hierarchical clustering (with different distance metrics; n=21 runs per EPG) or K-means clustering (3 different K; n=50 runs per EPG). Upper panel BRCA, lower Panel CRC. Violin plot distribution with median (blue square) and IQR.

We applied this approach to expression data sets for breast cancer (BRCA) and colorectal cancer (CRC). Clusters of gene co-expression were derived from resulting matrices using three approaches: hierarchical clustering, K-means clustering or a computationally efficient network tool, fast unfolding of communities in large networks (FastUnfold) [19]. In each instance clusters were generated from matrices in which genes retained all edges or compared to parsimonious correlation matrices retaining 3 to 10 edges per gene (Table S1). The resulting clusters (subsequently referred to as modules, Table S2) of co-expression were then tested for the separation of known biology, based on enrichment of ontology and signature terms. This was assessed as summative enrichment across signature terms and purity of enrichment, examining relative separation of biology between modules (Fig. 1C). The network method (FastUnfold) provided the most significant enrichment and segregation of ontology terms. Edge reduction improved the segregation of biology, and increasingly stringent edge-reduction enhanced the enrichment of ontologies/signatures and purity of segregation between modules across both cancers (Fig. 1C). Indeed, there was no significant benefit to retaining more than 3 edges per gene (EPG3). We call the EPG3 matrix clustered with FastUnfold a parsimonious gene correlation network analysis (PGCNA). Robustness of clustering was tested using the top 100 PGCNA clusterings, showing that for each cancer type modules retained a high proportion of the same genes across different clustering runs (Fig. S1B, C). For each cancer type the optimal PGCNA clustering based on ontology enrichment was taken forward for further analysis. Initially the networks were visualized as an interactive web-based resource (Fig. 2). To enhance network utility additional factors were overlaid providing inter-related visualizations of the data viewed through the networks (Fig. S2 & http://pgcna.gets-it.net/).

**Figure 2.**
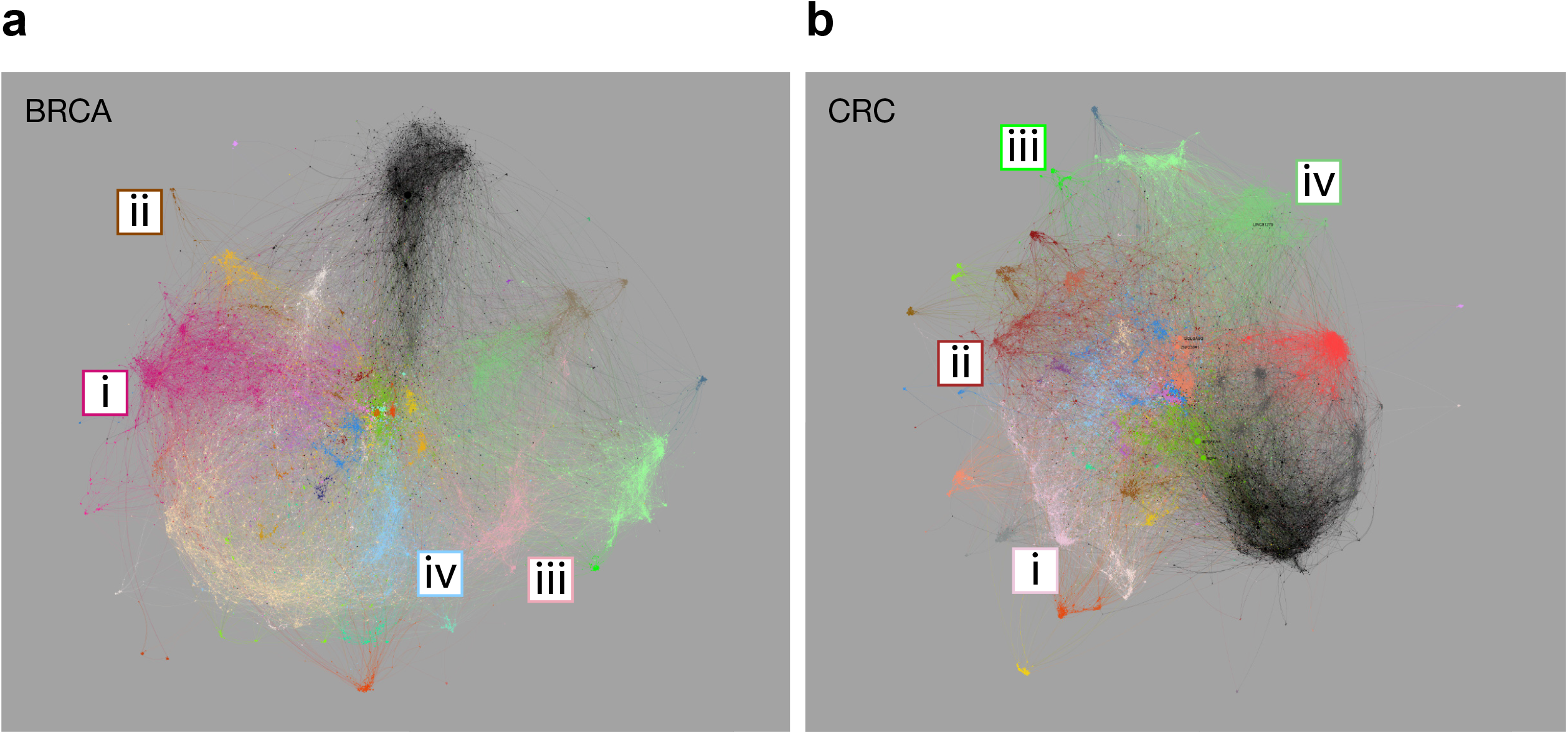
PGCNA network visualization for BRCA and CRC. **(A)** BCRA network with modules color-coded, modules overlapping significantly with those in CRC share a common color. Modules corresponding to intrinsic BRCA classification (i) luminal (BRCA_M6), (ii) ERBB2/HER2 (BRCA_M5), (iii) basal/normal (BRCA_M14), (iv) cell cycle (BRCA_M7). **(B)** CRC network, highlighted modules correspond to consensus molecular subtypes of CRC (i) CMS2-enterocyte (CRC_M3), (ii) CMS3-metabolic/goblet (CRC_M7), (iii) CMS1-hypermutated (CRC_M32), (iv) CMS4-mesenchymal (CRC_M8). Fully annotated versions in Fig. S2, and http://pgcna.gets-it.net/.

### Biology of network modules and mapping to expression-based cancer classifications

Detection of BRCA intrinsic sub-classes has been refined into expression-based tools such as the PAM50 classifier [5, 4, 8]. Mapping genes linked to these intrinsic classes onto the network identifies BRCA_M6 as the luminal module (Fig. 2A [i], S2A-C, online). Genes associated with ERBB2 amplified breast cancer map on to BRC_M5 (Fig. 2A [ii]), epithelial genes defining basal breast cancer overlap with those linked to normal-like breast cancers and map onto module BRCA_M14 (Fig. 2A [iii]). While genes linked to cell proliferation which provide a shared feature of Luminal B and Basal type breast cancers map onto BRCA_M7 (Fig. 2A [iv]).

The CRC consensus molecular subtype classification recognizes four subtypes [6]: CMS2 containing genes linked to canonical enterocyte-like differentiation maps onto module CRC_M3 (Fig. 2B [i]); CMS3 reflects goblet-cell and metabolic differentiation and maps onto CRC_M7 (Fig. 2B [ii]); CMS1 identifies microsatellite unstable cancers through interferon response genes and maps onto CRC_M32 (Fig. 2B [iii]); and CMS4 encompassing mesenchymal dominant CRC maps onto CRC_M8 (Fig. 2B [iv]) (Fig. S2D-F, online). Therefore, the PGCNA networks successfully place current paradigms of expression-based classification in BRCA and CRC in the context of wider expression patterns for each cancer.

Assessment of network clustering success was based on the enrichment and segregation of gene signatures between the resulting modules. These enrichments (Table S3, S4) were summarized to illustrate the most significantly enriched ontology and signature terms between modules (Fig. 3A, B & S3A, B). Purity of segregated biology was reflected in the separation of enriched signatures between individual modules. A summary designation was assigned to each module based on a selectively enriched term.

**Figure 3.**
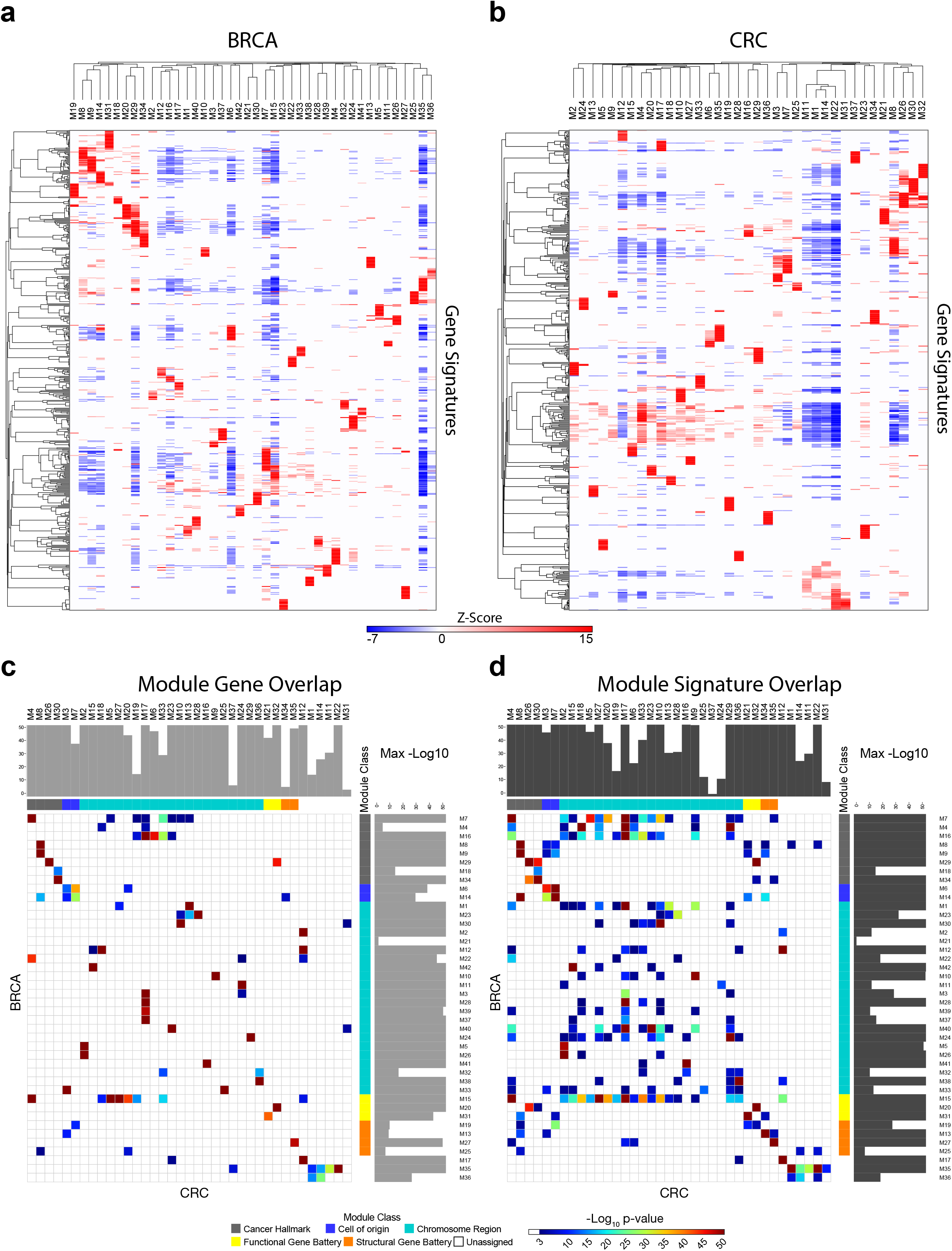
Module biology and between cancer analysis identifies principles of gene co-expression. Heatmaps of gene signature enrichment between modules **(A)** BRCA, **(B)** CRC. Significant enrichment or depletion illustrated on red/blue scale, x-axis (modules) and y-axis (signatures). Hierarchical clustering according to gene signature enrichment (using top 15 signatures per module; FDR <0.05). Scalable version in Fig. S3. **(C)** & **(D)** Module relationship between cancers analyzed using hypergeometric test displayed as pairwise comparison matrix. Significance of overlap displayed as p-values on indicated color scale (*p*-values < 0.001); overlap by **(C)** module gene membership, **(D)** enriched gene signatures. Grey side bars illustrate maximal significance for module match. Module class Cancer-Hallmark: gray, Cell-of-origin: blue, Chromosome-Region: cyan, Functional-Gene-Battery: yellow, Structural-Gene-Battery: orange and Unassigned: white.

We next tested whether recurrent features of cancer biology could be identified in the comparison of modules between the cancer types. Pairwise comparison demonstrated a high degree of similarity at the level of module gene membership (Fig. 3C, S3C) and ontologies/signatures associated with each module (Fig. 3D, S3D).

Considering cancer hallmarks recurrent modules could be identified relating to pathways linked to cell cycle, immune response, EMT/stroma and angiogenesis. Additional recurrent modules were linked to co-regulated gene batteries such as the IFN-response or growth factor signaling pathways, or structural gene clusters such as Histone, HOX and immunoglobulin genes. Moreover, these modules exhibited shared enrichments for signatures of transcription factor motifs linked to gene promoters (Table S5) [20]. In BRCA the impact of chromosomal copy number variation on gene expression in cis has been extensively analyzed [21]. Such patterns of gene co-expression were recovered in the networks and proved highly reproducible between BRCA and CRC, with the majority of BRCA modules linked to specific chromosomal region having a direct counterpart in CRC (Fig. 3 C, D).

Hence, the comparison between cancer types identified principle determinants of gene co-expression patterns. These reflect the impact of cancer hallmarks, functional and structural gene batteries, and copy number variation, which are overlaid on modules linked to the specific biology of the originating cell type.

### Module neighborhoods link to epithelial differentiation pathways

Within the individual modules, the network sub-structure identifies genes with the highest degrees of correlation. To resolve whether these patterns linked to discrete cell states we reran FastUnfold and signature enrichment analysis for modules independently and defined the resulting sub-structure as module neighborhoods (Fig. 4, Fig. S4, Table S6 & online).

**Figure 4.**
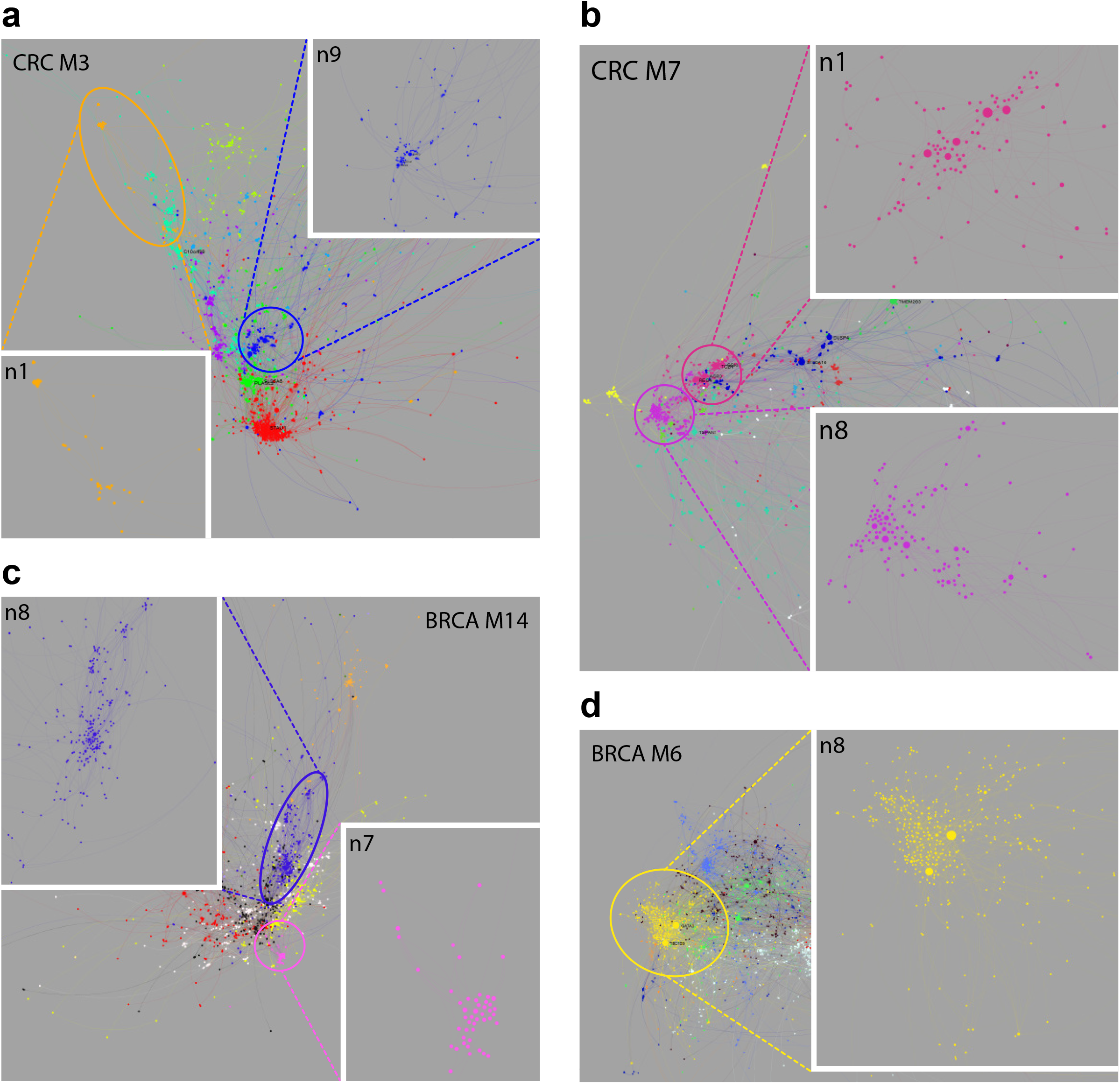
Module neighborhoods provide fine-grained resolution. Neighborhoods within modules are displayed by color code, interactive version online. **(A)** CRC_M3, enterocyte module, expanded: CRC_M3.n9, WNT-signaling (blue), and CRC_M3.n1, superficial enterocyte (orange). **(B)** CRC_M7, goblet metabolic module, expanded: CRC_M7.n8, classical goblet cell and CMS3 genes (purple), and CRC_M7.n1 putative deep secretory cell neighborhood (dark pink). **(C)** BRCA_M14, basal/normal module, expanded: BRCA_M14.n8 (blue), basal classifier genes, and BRCA_M14.n7 (pink), epithelial/epidermal differentiation. **(D)** BRCA_M6, luminal module, expanded: BRCA_M6.n8, *GATA3* and *ESR1* neighborhood (yellow).

In CRC features of epithelial differentiation are encompassed in CRC_M3 (enterocyte) and CRC_M7 (goblet cell). The enterocyte module encompassed neighborhoods enriched for genes linked to the WNT-signaling pathway (neighborhood 9, CRC_M3.n9), including *LGR5* [22], through to neighborhood CRC_M3.n1 enriched for genes characteristic of the mature enterocyte state (*CA1, CA4, CD177, MS4A12* and *SLC26A3*), recapitulating co-expression observed in single cell analysis of colonic epithelium (Fig. 4A) [23]. The goblet cell module divides into 11 neighborhoods of which 5 could be assigned to known ontology associations, for example CRC_M7.n10 linked to glycolysis and glucose metabolism and CRC_M7.n11 linked to defense responses (Fig. 4B, S4B). Neighborhood CRC_M7.n8, lacking enrichment of established ontology terms, included the hub genes *FCGBP* and *ST6GALNAC1* as well as *SPINK4* and *MUC2*, that are characteristic goblet cell markers linked to CMS3 CRC [23, 6]. The closely linked neighborhood CRC_M7.n1 included hub genes *REG4*, *AGR2* and *AGR3* (Fig. 4B, online). Notably REG4 has recently identified as a marker of deep crypt secretory cells [24].

In BRCA the luminal module (BRCA_M6) divides into 9 neighborhoods. Of these BRCA_M6.n8 is enriched for core ESR1 target genes and encompasses *GATA3* and *ESR1* as hub nodes (Fig. 4D, Table S6, online) [25, 26]. Genes that contribute to a basal-like classification and to epithelial biology fall in BRCA_M14. BRCA_M14.n8 includes the hub gene *SFRP1* as well as *EGFR* and *FOXC1*, PAM50 classifier genes used to define basal breast cancer (Fig. 4C, Table S6, online). A subset of basal breast cancer classifier genes are connected to the cytokeratin gene *KRT17* in BRCA_M14.n7 encompassing genes associated with epithelial and epidermal differentiation and linked to normal-like breast cancer classification (Fig. 4C, Table S6). Thus, the structure of gene neighborhoods in the epithelial modules reflects patterns of gene expression observed in differentiation, in both CRC and BRCA.

### Networks as multi-layered tools to explore survival associations

To provide resources that explore associations of expression with survival, we overlaid meta-information including association of gene expression with hazard ratio (HR) of death (Fig. 5, Fig. S5, online). The integration of multiple data sources retained the ability to detect robust HR associations. In the BRCA network, considered without histological subdivision, this recovered the separation of good and adverse outcome between luminal (BRCA_M6) and basal type (BRCA_M14) gene expression (Fig. 5A, S5A). At a module level cell cycle (BRCA_M7) showed the strongest adverse outcome association, which was also evident for modules linked to amplified chromosomal regions that cluster with the cell cycle module (such as BRCA_M24 & M37). Heterogeneity in HR association of module genes, as shown by spread in the violin plot across the neutral line, is a particular feature of the stem cell/EMT (BRCA_M9) and immune response modules (BRCA_M29). The latter encompasses several distinct neighborhoods (Fig. S5B), reproducing the ability to impute immune cell populations in cancer immune response from gene expression data. These separate into good outcome associations centered on T/NK- and B-lymphocyte genes or components of MHC class II, as opposed to genes characteristic of monocyte/macrophage populations such as *SLAMF8* and *CD163* link to adverse outcome.

**Figure 5.**
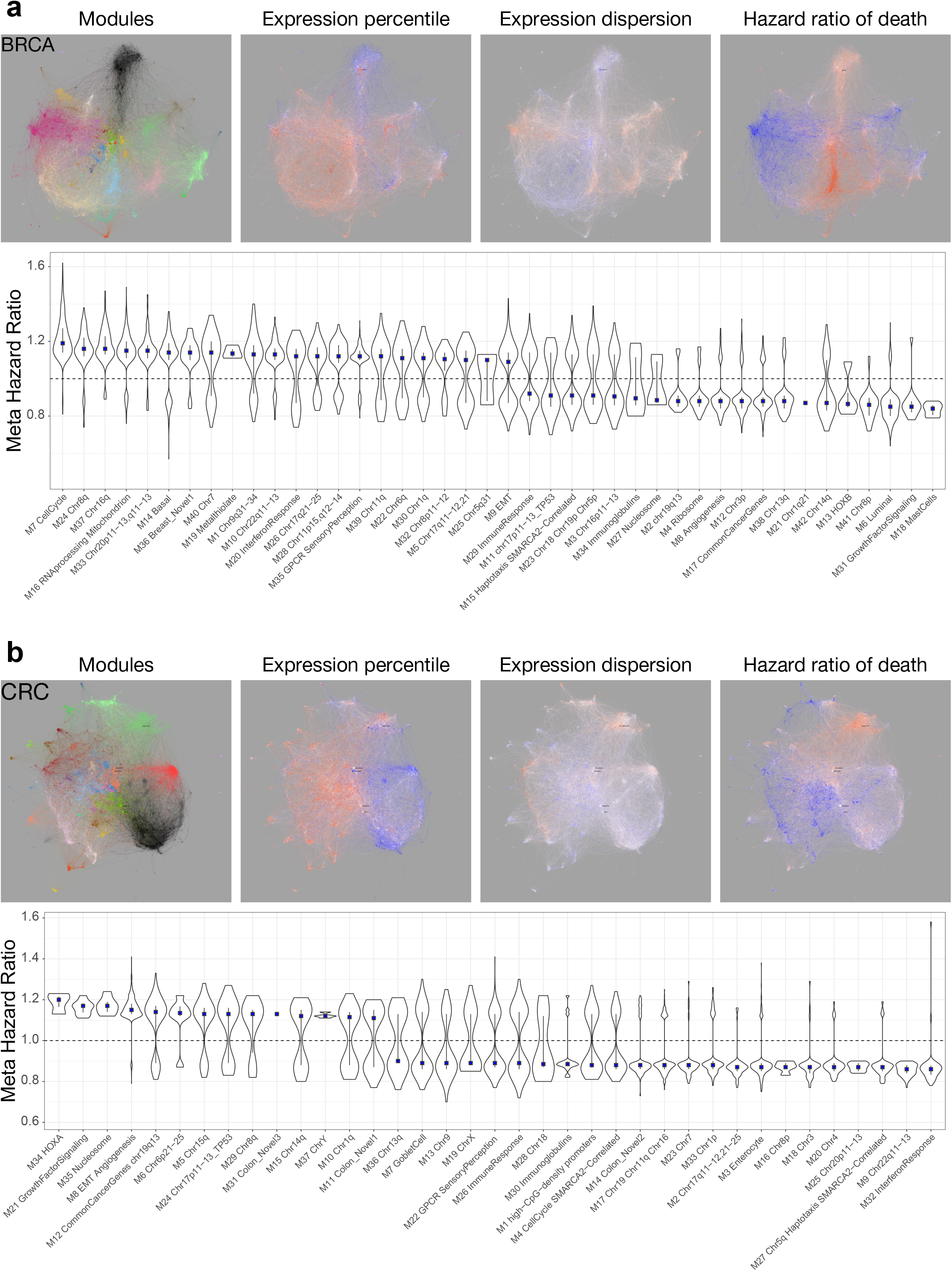
Networks as multilayered tools to explore survival association. BRCA **(A)** & CRC **(B)** meta-information overlay. Left to right: module color code, median expression percentile (relative intensity of expression) across data sets, median expression dispersion (Quartile coefficient of dispersion, variation between samples/patients) within data sets, and association of gene expression with meta-analysis HR (HR) of death. Color scales: expression dispersion and variance blue (least) to red (most); outcome blue (low HR - good outcome) to red (high HR - poor outcome). Lower panels, ranked module level association with HR of death. Distribution of HR associations for module genes with *p*-value < 0.05, along with median (blue square) and IQR.

In CRC, the enterocyte (CRC_M3) and interferon response modules (CRC_M32) were linked to good outcome, while adverse outcome associations centered on the EMT/angiogenesis module (CRC_M8) and modules linked to specific chromosomal regions (Fig. 5B, S5C). The three modules with strongest adverse outcome association were HOXA (CRC_M34), growth factor signaling (CRC_M21) and nucleosome (CRC_M35). In CRC the immune response module (CRC_M26) also showed a heterogenous pattern, distinct from the near homogenous good outcome association of the IFN module. Poor outcome associated with genes linked to macrophage/monocyte populations and again contrasted with good outcome for B- and T/NK-cell linked gene expression (Fig. S5D). Consistent with previous analysis [13], the immunoglobulin modules (BRCA_M34 & CRC_M30) indicative of tumor associated plasma cells were linked to good outcome, but with a relatively stronger signal in CRC.

### Network modules provide a platform for molecular stratification

Having validated PGCNA as a tool to interrogate the integrated training data sets, we next tested the modules as a platform to explore TCGA data [7, 27]. First, we used the 25 most representative genes (nodes) of each module to generate module expression values (MEV) and assessed module co-expression by hierarchical clustering. In both BRCA and CRC the overall pattern of module co-expression in RNA-seq data was closely related to that in array derived training data sets (Fig. S6) supporting the use of MEVs as a platform for analysis of TCGA data.

Applying the MEVs in hierarchical clustering segregated BRCA, initially without considering histological type, into branches according to expression of basal, luminal and mesenchymal related modules. In the latter this distinguished subsets or mesenchymal from mixed mesenchymal/angiogenic BRCA that included the majority of lobular breast cancers (Fig. S7A). Within these major branches further heterogeneity was evident across other network modules, sub-dividing the primary branches according to wider patterns of modular gene expression. Such subdivision was also evident within histological types (Fig. S8A) and for example illustrated a distinctive pattern of MEV expression in mucinous carcinomas.

Extending this approach to CRC the clustering divided into three main branches (Fig. S7B) corresponding to the primary features of the consensus molecular subtypes. However, the network modules again illustrated heterogeneity within these primary branches. This remained evident after separation by mutational load. Notably a subset of highly mutated CRC was identifiable as deficient in immune and EMT/angiogenesis module expression (Fig. S8B).

Thus, MEVs can be employed to capture both primary features related to existing consensus/intrinsic classes alongside features of gene expression across the wider characteristics of a cancer type.

### Network modules show distinctive mutational associations in BRCA and CRC

To integrate module expression with gene mutation we first considered BRCA as a single entity. This demonstrated a primary division of enrichment or anti-enrichment of *TP53* versus *CDH1*, *PIK3CA*, *GATA3*, *MAP3K1*, *KMT2C* and *NCOR1* mutation (Fig. 6A). *TP53* mutation positively correlated with the cell cycle and basal modules, and some chromosomal regional modules. Cell cycle as well as the immune response and IFN modules were additionally distinguished by a positive association with diverse additional mutational targets. The luminal, EMT, angiogenesis and related modules were significantly anti-correlated with *TP53* mutation and positively associated with combinations of mutations in *CDH1*, *PIK3CA*, *GATA3*, *MAP3K1*, *KMT2C* and *NCOR1*.

**Figure 6.**
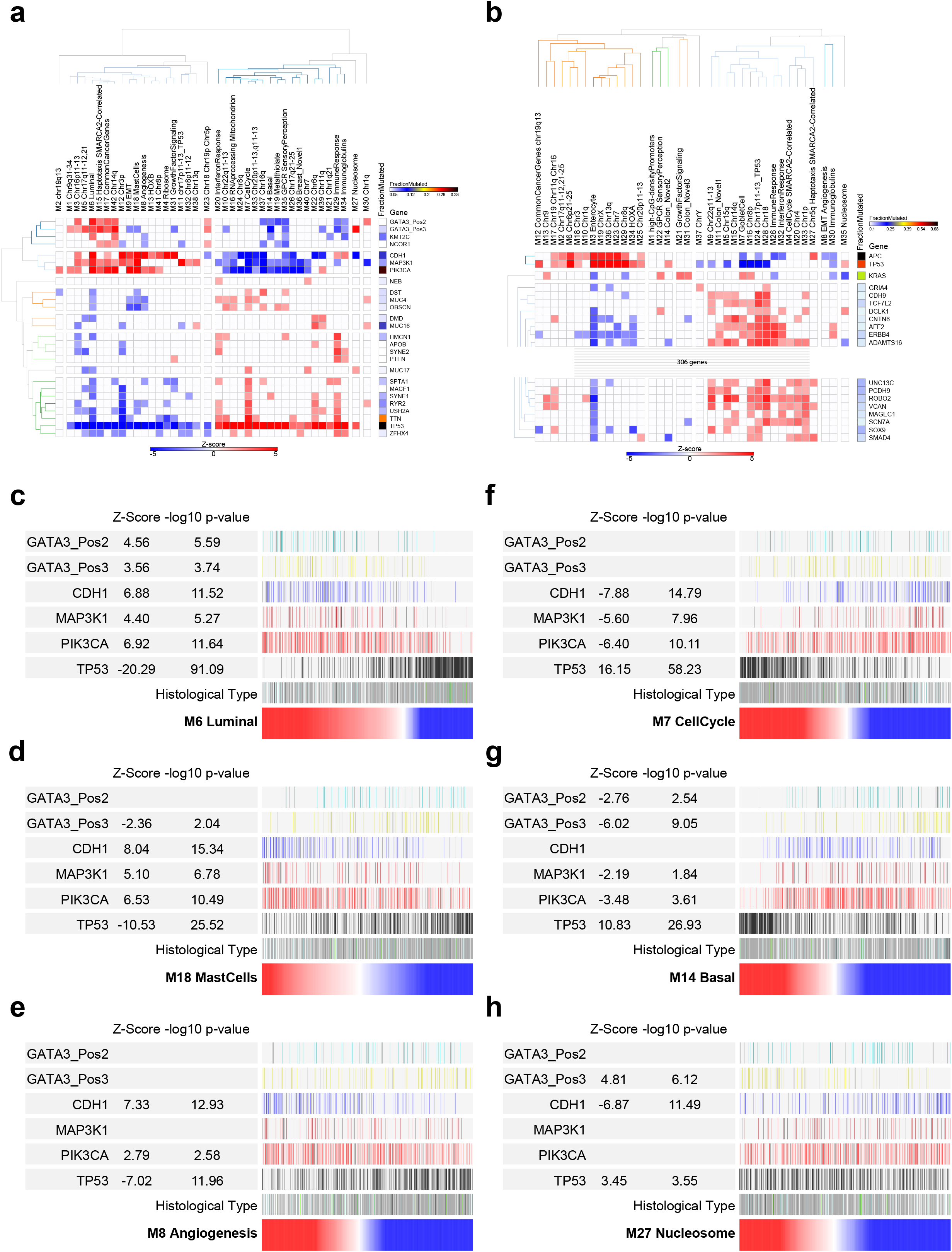
Network modules show distinctive patterns of mutational association. Correlations of MEV with mutation status of commonly mutated genes in TCGA data **(A)** BRCA & **(B)** CRC. Significance of Spearman’s Rank correlation of MEV with mutation, z-score on blue to red scale; fraction of mutated cases per gene, blue to black color scale. Hierarchical clustering for genes mutated in ≥ 5% BRCA & ≥ 10% CRC of TCGA samples. For CRC the heatmap is truncated for display purposes (complete version Fig. S9C). **(C)** BRCA_M6 Luminal, **(D)** BRCA_M18 mast cell, **(E)** BRCA_M8 angiogenesis, **(F)** BRCA_M7 cell cycle, **(G)** BRCA_M14 basal, and **(H)** BRCA_M27 nucleosome, MEVs as ranking variables (red to blue color scale) for mutation distribution, z-score and −log10 p-value for *GATA3* with division into proximal (pos2;N-terminus) and distal (pos3; C-terminus), *CDH1*, *MAP3K1*, *PIK3CA* and *TP53*. Histological: ductal (grey), lobular (white) lobular/ductal (dark blue), medullary (green), metaplastic (dark green), mucinous (black), not reported (light blue).

*GATA3* mutations can be subdivided between DNA-binding domain or carboxy-terminus, with the latter including frameshift mutations. The potential value of MEVs as a tool for assessing selective patterns of gene expression is supported by the observation that *GATA3* mutations affecting the carboxy-terminus are selectively associated with nucleosome module expression. Extending the analysis to BRCA after division by histological subtype, the general pattern observed in all BRCA irrespective of histological type was evident when considering ductal BRCA in isolation (Fig. S9A). Lobular BRCA is more molecularly homogenous, reflected in a sparse correlation pattern, nonetheless also retaining features observed in BRCA as a whole (Fig. S9B).

For CRC, the pattern was impacted by the high overall mutation load (Fig. 6B, S9C). Associations divided around modules linked with both *TP53* and *APC* mutation and those that correlated with a high mutation load across a wide range of target genes and that were neutral or anti-correlated with *TP53* and *APC* mutation. This separated the enterocyte module, linked to *TP53* and *APC* mutation, and the goblet cell module linked to high mutation load and *KRAS* mutation, with *KRAS* mutation also correlating with the growth factor signaling module. In CRC the cell cycle module was not positively correlated with *TP53* mutation, but instead was linked to the broad swathe of highly mutated target genes. The modules most strongly linked to mutation load encompassed genes from the vicinity of the *TP53* on chr17p, chr18 and components of the immune response and IFN signaling. Overall this reinforces the division of CRC into the major molecular pathways of *TP53* and *APC* mutation versus hypermutational genomic instability and supports the broadly different patterns of molecular features linked to patterns of goblet cell or enterocyte module expression in CRC.

### Patterns of mutation link to intensity of module expression

Finally, we addressed the potential value of module expression intensity. The BRCA luminal MEV intensity showed a strong positive correlation with *CDH1*, *MAP3K1*, *GATA3* and *PIK3CA* mutations and profound anti-correlation with *TP53* mutation (Fig. 6C). This was paralleled by the opposite association of cell cycle (Fig. 6F) and basal (Fig. 6G) MEV intensity with these mutations. The angiogenesis module separated a strong positive association of expression intensity with *CDH1* mutation status from either *GATA3* or *MAP3K1* mutation. By contrast the mast cell MEV outperformed the luminal module as a ranking variable in relation to *CDH*1 and *MAP3K1* mutation across both lobular and ductal type BRCA (Fig. 6D), which was notable given the link between mast cell module and good outcome. Contrasting with this was the nucleosome module which when used as a ranking variable emphasized selective positive correlation with *GATA3* 3’-mutations (Fig. 6H), and anti-correlation with *CDH1* mutation status. As part of this specific association, both nucleosome module expression and *GATA3* 3’ mutations were enriched in mucinous BRCA (p-value 0.0004). We conclude that use of MEVs as ranking variables illustrated a principle that the extremes of module expression selected for increasingly stereotyped tumors with more distinct patterns of mutation association.

## Discussion

We set out to test whether the modular nature of gene co-expression could be used to derive expression codes summarizing diverse features of cancer biology. Furthermore, whether these could enhance molecular stratification by providing a link between existing assets of large gene expression data and resources of multi-parameter cancer exploration exemplified by TCGA.

A striking finding is that radical pruning of edges in expression correlation matrices prior to network analysis with FastUnfold allows remarkably efficient recovery of biology. Neighborhoods of highly correlated genes within network modules recover patterns of gene co-expression observed in previous single cell analysis of cellular sub-populations [23]. The patterns of association seen at the gene neighborhood level typified by the immune response modules recapitulate features seen in previous analyses [13, 12], while allowing extension to the single gene level. Thus, the network and its modular structure may be used at different levels to separate or coalesce cellular features.

CRC and BRCA show a remarkable communality in gene co-expression patterns. The shared biology supports a set of core principles that underpin patterns of co-expression in cancer. These can be summarized as (1) genes linked to cancer hallmark features such as cell cycle, EMT, angiogenesis, and immune response; (2) functional gene batteries linked to either specific pathways such as the IFN-response or growth factor receptor signaling or to structural clusters of co-regulated genes; and (3) to co-expression related to copy number variation. In each case these shared drivers are overlaid on modules derived from the selective biology of the originating lineage.

As a platform from which to enrich molecular stratification, the networks recover modules that map closely onto existing classifications for both BRCA and CRC, and place these in a wider context. Using hub genes to generate MEVs allowed the integration of expression with mutation profiles in the TCGA resource at data set and case-by-case level, in effect exploring the TCGA data from the perspective of the deep expression data available for BRCA and CRC. Together these provide evidence that molecular classification may be enriched by using MEVs as a gene expression barcode, complementing current paradigms.

The approach we describe here has both disease-specific and general relevance. It provides an approach for extracting useful networks that can be applied effectively to diverse clinical and experimental data sets, while also generating a mineable resource, and illustrates how resulting network modules might be used to sit alongside existing expression-based classifications to enhance molecular stratification.

## Acknowledgements

This work was supported by Cancer Research UK programme grant (C7845/A17723). We thank Gina Doody and Ulf Klein for critical review of the manuscript.

## Author contribution

MAC performed analyses, analyzed results, conceived study, wrote paper, DRW guided analyses, edited paper, RMT conceived study, analyzed results, wrote paper.

## Methods

### CONTACT FOR REAGENT AND RESOURCE SHARING

Further information and requests for resources and reagents should be directed to Matthew Care or Reuben Tooze.

### METHOD DETAILS

See Supplemental Figure 1 (Fig. S1A) for outline, will refer to numbers in this figure in the sections below.

#### Expression data sets

See Fig. S1A part 1

For the generation of the gene correlation networks 23 breast cancer (BRCA) and 12 colorectal cancer (CRC) gene expression data sets were downloaded from the Gene Expression Omnibus(Barrett et al., 2011) (See Key Resources Table) (BRCA, 7464 cases; 26 arrays) [28-49] and (CRC, 2399 cases; 11 arrays after merging of 2)[50-59]. Three of the BRCA data sets were on two different expression platforms (GSE3494, GSE36774 and GSE4922), these were analyzed independently, giving a total of 26 BRCA expression data sets. In the case of CRC two related data sets were merged (GSE17536), giving a total of 11 CRC data sets.

#### TCGA data sets

For independent assessment of the network modules two RNA-seq data sets were downloaded from The Cancer Genome Atlas (BRCA/CRC data sets were downloaded on 2017.11.15 from http://cancergenome.nih.gov/) along with the corresponding simple nucleotide variation data (MuTect2 pipeline). The overlapping expression/mutation samples were used for downstream analyses (See Key Resources Table).

Normalization and re-annotation of data

For each data set the probes were re-annotated using the MyGene.info (http://mygene.info) API using all available references (e.g. NCBI Entrez, Ensembl etc.) and any ambiguous mappings manually assigned [60].

Each data set was quantile normalized using the R Limma package and the probes for each gene merged by taking the median value for probe sets with a Pearson correlation ≥0.2 and the maximum value for these with a correlation <0.2 [61].

#### Network analysis

This discusses how the Parsimonious Gene Correlation Network Analysis (PGCNA) approach was developed.

#### Gene correlation calculation

See Fig. S1A part 2 and 3

For each expression data set the 80% most variant genes were used to calculate Spearman’s rank correlations for all gene pairs using the Python scipy.stats package. The resultant *p*-values and correlations matrices were merged across all data sets for a given cancer by taking the median values (across the sets in which the gene pairs were contained) to give a final median correlation matrix and its corresponding *p*-value matrix. Genes present in < 9 data sets for BRCA and < 4 data sets for CRC were removed from respective matrices. This gave a final matrix size of 17,805 and 18,896 for BRCA and CRC respectively. Finally, all correlations with a *p*-value > 0.05 were set to 0 to reduce noise.

#### Edge reduction

See Fig. S1A part 4

We tested a simple but aggressive edge reduction strategy as a way to improve module discovery and network visualization. For each gene (row) in a correlation matrix only the N most correlated Edges Per Gene (EPG) were retained, with N ranging from 3 to 10 (<3 gives orphan modules). The resulting matrix *M*, with entries written as *M* = (*m_ij_*) was made symmetrical by setting *m_ij_ = m_ji_* for all indices *i* and *j* so that *M* = M^T^ (its transpose). For EPG3 this reduced the nodes in BRCA from 43,231,589 to 49,199 and CRC from 42,142,502 to 52,257, in both cases > 800-fold reduction (Supplemental Table 1).

#### Data clustering

See Fig. S1A part 5

The matrices from the edge reduction step alongside the Total matrices were clustered using 3 different approaches: hierarchical clustering using the R package fastcluster, k-means clustering using the R package kmeans and a network level clustering using the Fast unfolding of communities in large networks algorithm (version 0.3) referred to subsequently herein as FastUnfold [19, 62]. FastUnfold was run 10,000 times at each EPG level and the 100 best (judged by the modularity score) were used for downstream analysis (note: the Total edges data always yielded 3 modules and was thus ignored for the FastUnfold approach). The FastUnfold algorithm automatically converges on a module number and therefore does not require a user defined module number.

For the k-means clustering k was set to ± 1 around the module number from the best FastUnfold solution (see Cluster selection) and for each k and EPG 50 iterations were run. For hierarchical clustering 8 different linkage methods (average, centroid, complete, Mcquitty, Median, Single, WardD and WardD2) were used and the resultant dendrograms cut at ± 10 around the module number from the best FastUnfold solution giving 21 results for every input matrix (note: only the 2 best linkage methods, WardD/WardD2, are shown in Figure 1C).

#### Computational efficiency

See Fig. S1A part 5

All data clustering was run on the MARC1 HPC at the University of Leeds. For comparison of computational efficiency (averaged between BRCA/CRC): For FastUnfold EPG3 after 47 seconds initial setup 50 data clustering ran in 10 seconds with memory usage of 60 MB.

For k-means for 50 clusterings of the total data the run time was ∼30h with memory usage of 25GB. While for EPG3 the run time was ∼15h with memory usage of 24.8GB. For hierarchical clustering the total run time was ∼6.5h with memory usage of ∼16GB for both total and EPG3.

#### Cluster selection

See Fig. S1A part 6

The success of the clustering approaches was assessed by looking at the level of biological enrichment of each module while rewarding purity (biological enrichment in single modules) and similar (even) module sizes (i.e. to avoid skewing to a few modules that contain many genes/functions).

Gene signature analysis was carried out for each module, from each clustering of the data. Then to generate a total enrichment score for a given clustering:

Signatures were filtered to retain only those with ≥ 5 and ≤ 1000 genes with an FDR (Benjamini Hochberg) of < 0.05.

For each module within a clustering, the enriched signatures were ranked by FDR and the top 15 added to a global list of signatures for that clustering.

A matrix was generated that contained all the z-scores for every signature (rows) in the global list across all the modules (columns).

For each signature a fractional contribution was calculated as the row-max-zScore/row-sum-zScores (where 1 = enrichment of signature in only 1 module). Across all signatures a median factional contribution (MFC) was calculated.

The sum of the maximum z-score per signature (row) was calculated (ZScoreMS). Module size skewing was assessed by calculating the normalized Shannon entropy:

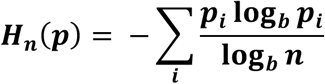

of the module sizes. This gave a score that ranged from 1 (even module sizes) towards 0 with increasing skewing.

A final clustering enrichment score was calculated as: ZScoreMS ⋅ MFC ⋅ normalizedEntropy.

This allowed the selection of the best FastUnfold clustering (Figure 1C; Gene Signature Enrichment: FastUnfold). This was then used to set the module number range in the k-means/hierarchical approaches. The FastUnfold method outperformed the k-means/hierarchical clustering methods across all EPG, with only the Ward-linkage hierarchical clustering approaching a similar enrichment when using the Total data. With increasing EPG there was a corresponding decrease in module number with no trade-off of increased biological enrichment (Figure 1B and 1C). Thus, for all downstream analysis we chose the optimal FastUnfold EPG3 result for both cancers; this combination of FastUnfold and EPG3 we term a Parsimonious Gene Correlation Network Analysis (PGCNA). However, it should be noted that most of the recovered modules were broadly retained across the 100 FastUnfold clustering results (see Module Stability and Supplemental Figure 1C/D).

Figure 3A and Supplemental Figure 3A/B show visualizations of the optimal BRCA/CRC gene signature results. As before these show the top 15 signatures per module (with ≥ 5 and ≤ 1000 genes) but are filtered with the more lenient *p*-value < 0.01.

#### Module stability

The stability of modules was assessed to see how recurrent the modules were across different clustering runs (Supplemental Figure 1B/C). Using the optimal clustering as a reference, for each of the 100 FastUnfold clustering, per reference module:

Find the maximum overlapping module.

Store the number of overlapping genes along with significance (*p*-value) of the overlap and increment sums for the overlapping genes.

The stability % per gene is simply the overlap sum (i.e. across 100 clustering runs what % of maximum overlapping modules is the gene found in). The stability values per reference module was calculated as median overlap across the 100 clustering runs.

#### Network visualization

See Fig. S1A part 7

The optimal EPG3 matrix from BRCA/CRC was converted into a list of edges and nodes and uploaded into the Gephi package (version 0.82) [63]. Modules were colored so that where possible significantly overlapping modules between BRCA and CRC shared colors. Degree and Betweenness Centrality were calculated and the latter used to adjust node sizes. The network layout was generated using the ForceAtlas2 approach [64], and interactive HTML5 web visualizations exported using the sigma.js library (https://github.com/oxfordinternetinstitute/gephi-plugins/tree/sigmaexporter-plugin).

#### Network meta-data

See Fig. S1A part 8

A number of additional features were calculated for the network genes across the data sets used to generate the correlation network. For each gene the median percentile expression was calculated across all data sets, its dispersion across data-sets calculated as the median absolute deviation (MAD) and its dispersion within data-sets (i.e. across patients) calculated as the median quantile coefficient of dispersion (QCOD).

The Survival library for R was used to analyze right-censored survival data for the data sets where this was available (n=8 for BRCA, n=4 for CRC). Within each data set the expression of each gene (as z-score) was used as a continuous variable in a Cox Proportional Hazards model. Across data sets a meta-analysis was conducted by fitting a fixed-effect model (R metafor package) to the hazard ratios, weighted by data set size. rma(yi=lnHazardRatio,sei=standardErr,weights=dataSetSize,weighted=TRUE,method=" FE") [65].

#### Module overlaps

See Fig. S1A part 9

The overlap of the modules between that cancers at the gene and signature level was assessed using a hypergeometric test and the overlap visualized as a python matplotlib heatmap of −log_10_ *p-*values (Fig. 3B & C), and with the overlap number and module size displayed (Fig. S3C & D). The signatures were pre-filtered to *p*-value <0.001 and ≥ 5 and ≤ 1000 genes.

#### Application to TCGA data

The modules derived from the GEO ‘training data’ were used to analyze unseen expression and mutation data from The Cancer Genome Atlas (TCGA).

#### Module Expression Values

See Fig. S1A part 10

To assign module enrichment/depletion at the patient level a summary score was created for each module for each patient.

Within each data set, which vary in available genes, the first step was to select the 25 most representative genes per module: For every gene a connectivity score was calculated by summing its correlations within its module.

This was then weighted using expression and dispersion information

ModCon = connectivity^2^⋅ percentileExpression⋅VarWithin⋅(100 – VarAcross)/100

Where VarWithin is the dispersion of a gene expression within data sets measured as the median quantile coefficient of dispersion (max range 0—1), VarAcross is the dispersion of gene expression across data sets measured as the median absolute deviation of percentile expression (max range 0—100). This rewards genes that have high connectivity and are variant across patients but invariant across data sets.

Genes were ranked by ModCon and the top 25 selected.

These 25 genes were then converted to a Module Expression Value (MEV): Per gene, standardize (z-score) the quantile normalized log_2_ expression data. Per sample (patient) sum the 25 z-scores to give a MEV.

#### Heatmap visualizations

See Fig. S1A part 11

The MEV were used to create heatmap visualizations of each module at the patient level within the BRCA and CRC TCGA data sets. Using the Broad GENE-E package (https://software.broadinstitute.org/GENE-E/) the MEV were hierarchically clustered (Pearson correlations and average linkage) and displayed along with available meta data (Fig. 6 & Fig. S7).

#### Mutation correlation analysis

See Fig. S1A part 12

The relationship of mutations and modules was calculated using the MuTect2 simple nucleotide variation (SNV) mutation data and the MEV. The SNV data was filtered to retain mutations present in > 5 or > 10% of patients in BRCA and CRC respectively. Spearman’s rank correlations were calculated between all pairs of mutated gene and module. These were converted to z-scores to convey the ± correlation along with its significance. A matrix was output containing the z-scores for all gene/modules ≥ 1 positive significant (*p*-value <0.05) correlation (i.e. a gene need only be significant in one module to be included). This matrix was then hierarchically clustered (Pearson correlations and average linkage) using GENE-E (Figure 7 and Supplemental Figures 8). For BRCA the 140 GATA3 mutations were split into 3 groups based on mutation position: GATA3_Pos1 (Chr10: 8058419—8064131; n=10), GATA3_Pos2 (Chr10: 8069470— 8069596; n=57) and GATA3_Pos3 (Chr10: 8073734—8074229; n=73).

### QUANTIFICATION AND STATISTICAL ANALYSIS

#### Gene signature data and enrichment analysis

A data set of 17,211 gene signatures was created by merging signatures downloaded from http://lymphochip.nih.gov/signaturedb/ (SignatureDB), http://www.broadinstitute.org/gsea/msigdb/index.jsp MSigDB V6.1 (MSigDB C1–C7 and H; excluding C5. With MIPS signatures from version 3.1 and PID signatures from version 4 added back), http://compbio.dfci.harvard.edu/genesigdb/ Gene Signature Database V4 (GeneSigDB), UniProt keywords (parsed XML from http://www.uniprot.org/downloads), and locally curated lists. A gene ontology gene set was created using an in-house python script. This parses a gene association file (http://geneontology.org/page/download-go-annotations) to link genes with ontology terms and then uses the ontology structure (.obo file; http://purl.obolibrary.org/obo/go.obo) to propagate these terms up to the root. The resultant gene set contained 22,271 terms. The gene-ontology and gene-signatures sets were merged to give a final signature set of 39,482 terms.

Enrichment of gene lists for signatures was assessed using a hypergeometric test, in which the draw is the gene list genes, the successes are the signature genes, and the population is the genes present on the platform.

#### Correlation of modules

The relationship of the modules was analyzed by calculating the Spearman’s rank correlation for all module (as MEV) pairs within each data set. These were then merged across data sets by calculating the median correlation and *p*-values. A final matrix generated by setting all correlations with a *p*-value > 0.05 to 0. Within GENE-E the ‘training data’ was hierarchically clustered (Pearson correlations and average linkage) and the TCGA data displayed in the same order without hierarchical clustering (Supplemental Figure 6).

### DATA AND SOFTWARE AVAILABILITY

Interactive networks and all meta-data is available at http://pgcna.gets-it.net/. All scripts are available upon request

## Supplemental Figure Legends

**Figure S1. Process diagram and module stability. Accompanies Figure 1 (A).**

(A) A detailed version of the process diagram with numbering of the process linked to Online Methods sections. The diagram is divided into the broad categories of Development & Validation, Data Exploration and Network Testing. (B) BRCA and (C) CRC, violin plots of the stability of module membership generated in the 100 networks evaluated for EPG3 using fast unfolding networks. The module membership of the optimal clustering was set as reference. The degree of overlap of module gene membership with these reference modules is shown as percentage stability across the other 99 network clusterings. Violin plots display the distribution along with median (blue square) and the IQR, ordered by module median stability.

**Figure S2. Network visualization. Accompanies, Figure 2 and Figure 5.**

Detailed annotation of the network generated by PGCNA for BRCA displaying the optimal clustering as shown in Fig. 2, with all modules annotated and labelled with their respective summary designation. (A) displays primary module color coding for BRCA, with shared coding where applicable to the closest matching module for CRC in (D), (B) shows module annotations in the context of the network meta hazard ratio overlay, and (C) annotation in the context of expression percentile. (D) Annotated primary module color coded version of CRC network, (E) annotation in relation to meta hazard ratio, and (F) annotation in the context of expression percentile. Color scales: expression percentile blue (least) to red (most); outcome blue (low HR - good outcome) to red (high HR - poor outcome). Networks and linked resources are available as fully searchable interactive tools at at http://pgcna.gets-it.net/

Figure S3. Gene signature and ontology enrichments and overlap of module gene membership between BRCA and CRC. Accompanies Figure 3.

High-resolution version of heatmaps of gene signature and ontology term enrichments for the network modules of (A) BRCA and (B) CRC. Module numbers and designations are listed on the x-axis, and signature/ontology terms on the y-axis, modules were clustered according to gene signature enrichment using hierarchical clustering (using top 15 signatures per module; FDR <0.05). Gene signature enrichments are illustrated as a red/blue color code reflecting significance (z-score) of enrichment, complete lists of all signature enrichment results including the contributing genes are provided in SI-Table 3&4 and online resources. (C) Heatmap of module gene membership overlap and (D) module gene signature enrichment overlap in the pairwise comparison of BRCA (y-axis) and CRC (x-axis) modules. In each instance the number of genes included in the module is show along with the module number and summary term. Within the pairwise comparison the significance of overlap is illustrated in the indicated color scale of −-log10 p-value. For each pairwise comparison the number of overlapping genes is indicated in the relevant square.

**Figure S4. Gene signature and ontology enrichments in epithelial differentiation module neighborhoods. Accompanies Figure 4.**

These heatmaps illustrate the results of gene signature and ontology term enrichment analysis for the module neighborhood analysis shown in Figure 4, (A) CRC_M3 enterocyte, (B) CRC_M7 goblet cell, (C) BRCA_M6 luminal, (D) BRCA_M14 basal. Neighborhood numbers and designations are listed on the x-axis, and signature/ontology terms on the y-axis, modules were clustered according to gene signature enrichment using hierarchical clustering (using top 15 signatures per module; FDR <0.05). Gene signature enrichments are illustrated as a red/blue color code reflecting significance (z-score) of enrichment, complete lists of all signature enrichment results including the contributing genes are provided in SI-Table 6 and online resources.

**Figure S5. Hazard ratio overlays provide tools for analyzing potential prognostic associations. Accompanies Figure 5.**

This figure displays the relationship between hazard ratio at module and gene level for the BRCA and CRC networks. (A) Highlights the position of luminal, basal and cell cycle related modules that map onto components of the intrinsic classes of BRCA in relation to meta hazard ratio (lower panel). (B) Shows an expanded illustration of the immune response components of the BRCA network, with neighborhoods identified with indicative labels and showing the relationship to the relative position in the original color-coded network for comparison (left panel: meta hazard ratio, right panel: module colors). (C) Highlights the position of enterocyte, goblet and mesenchymal modules that map onto components of the consensus molecular subtype classes of CRC in relation to hazard ratio (lower panel). (D) Shows an expanded illustration of the immune response components of the CRC network, with neighborhoods identified with indicative labels and showing the relationship to the relative position in the original color-coded network for comparison (top panel: meta hazard ratio, bottom panel: module colors). (E) Illustrates the core components of the growth factor signaling modules of BRCA and CRC with the juxtaposition of the hazard ratio overlay (right panel in each case). Meta hazard ratios are shown on a color scale from blue (low) to red (high)

**Figure S6. Assessment of network module co-occurrence across all samples in training array data sets and TCGA RNA-seq data. Accompanies Figure 6.**

Correlation heatmaps of the co-occurrence of network modules in array training data (left panels) and TCGA RNAseq data (right panels) for BRCA (upper panels) and CRC (lower panels). Module expression values (MEV) were generated for all samples from the 25 (or less for smaller modules) hub genes of each module (see Online methods). The relationship of the modules was analyzed by calculating the Spearman’s rank correlation for all module (as MEV) pairs within each data set. These were then merged across data sets by calculating the median correlation and p-values. A final matrix generated by setting all correlations with a p-value > 0.05 to 0. Within GENE-E the ‘training data’ was hierarchically clustered (Pearson correlations and average linkage) and the TCGA data displayed in the same order without hierarchical clustering.

**Figure S7. Module expression values as a platform for a bar-code of gene expression**

Use of module expression values in hierarchical clustering of TGCA data (A) BRCA. Illustrated are the distribution of mutations in common target genes above the expression values as colored bars GATA3_Pos2 (green), GATA3_Pos3 (yellow), CDH1 (blue), MAP3K1 (dark red), PIK3CA (red), TP53 (black) as well as a wider range of less frequently mutated target genes (grey). Beneath this is illustrated assessment of highly mutated status and mutation burden, ER and PR status, and histological type. Beneath the heatmap indicative examples of the modules linked to heterogeneity in the principle branches of the tree are identified, as indicated in the figure key to the left of the heatmap. (B) CRC. Illustrated above the heatmap are the distribution of mutations in common target genes as colored bars, TP53 (black), KRAS (red) and APC (blue), beneath these mutation events in a wider set of representative genes (grey). The distribution of dichotomized highly mutated cases is shown in red, beneath this refined assessment of mutation burden is provided (blue to red color scale). A range of other meta-data is indicated in the figure key to the left of the heat-map and includes histological type, anatomical subdivision, pathological stage, colon polyps, BRAF gene analysis, micro-satellite instability. Beneath the heatmap indicative examples of the modules linked to heterogeneity in the principle branches of the tree are identified.

**Figure S8. Module expression values in stratification of BRCA after subdivision by histological type, and CRC after division by mutational load. Accompanies Figure 6.**

(A) Illustrates the results of applying hierarchical clustering using MEVSs for BRCA cases after separating by histological type. Illustrated are the distribution of mutations in common target genes above the expression values as colored bars GATA3_Pos2 (green), GATA3_Pos3 (yellow), CDH1 (blue), MAP3K1 (dark red), PIK3CA (red), TP53 (black) as well as a wider range of less frequently mutated target genes (grey). Beneath this is illustrated assessment of highly mutated status and mutation burden, ER and PR status, and histological type. Beneath the heatmap indicative examples of the modules linked to heterogeneity in the principle branches of the tree are identified, as indicated in the figure key to the left of the heatmap. (B) Illustrates the results of hierarchical clustering of CRC cases using MEVs after subdivision by hypermutation status. Illustrated above the heatmap are the distribution of mutations in common target genes as colored bars, TP53 (black), KRAS (red) and APC (blue), beneath these mutation events in a wider set of representative genes (grey). The distribution of dichotomized highly mutated cases is shown in red, beneath this refined assessment of mutation burden is provided (blue to red color scale). A range of other meta-data is indicated in the figure key to the left of the heat-map and includes histological type, anatomical subdivision, pathological stage, colon polyps, BRAF gene analysis, micro-satellite instability. Beneath the heatmap indicative examples of the modules linked to heterogeneity in the principle branches of the tree are identified.

**Figure S9. Network module and mutational association for BRCA divided by histological type and CRC. Accompanies Figure 7.**

This figure shows the results of analyzing the correlation between module expression values (MEV) and mutations in BRCA/CRC after separating cases according to histological type for (A) infiltrating ductal BRCA, (B) infiltrating lobular BRCA and (C) complete version of the correlations between module expression values (MEV) and mutation status for CRC, as shown in a truncated format in Fig. 6B. Significance of the Spearman’s Rank correlation of MEV with mutation is illustrated as a z-score with the indicated blue to red scale, while for each gene the fraction of mutated cases and mutation type are illustrated with blue to black color code along the side of the heatmap. Heatmap shows hierarchical clustering for genes mutated in ≥ 5% BRCA & ≥ 10% CRC of TCGA patients.

